# Small-molecule IGF1R inhibitors extend healthspan in a mouse model

**DOI:** 10.64898/2026.04.13.718278

**Authors:** Yulia Balandina, Tom Stadnikov, Gregory Basarab, Charles J. Eyermann, Alexander Suvorov

## Abstract

Antagonistic pleiotropy of the IGF-1 signaling cascade is well recognized, as it promotes growth and development at younger ages and delays aging later in life. The goal of this study is to test in a mouse longevity experiment whether orally delivered small-molecule IGF1R inhibitors have promise as an anti-aging therapy. C57BL/6 mice (25 male and 25 female mice per treatment) were treated with selective IGF1R inhibitors, picropodophyllin (PPP) or 5-[3-(phenylmethoxy)phenyl]-7-[trans-3-(1-pyrrolidinylmethyl)cyclobutyl]-7H-pyrrolo[3-d]pyrimidin-4-amine (NVP-ADW742), via powdered diets starting at 13 months of age, and physiological and behavioral parameters, as well as survival, were assessed. Both compounds protected both sexes from short-term memory decline; reduced systolic blood pressure in males and pulse rate in both sexes; rescued declining glucose tolerance in males; and abolished grey hair development, reduced frailty, and protected against decline in grip strength in female mice. There were no sex differences in survival curves within groups. No significant differences between groups were observed in the Kaplan–Meier analysis. However, the survival curve in the NVP-ADW742 group was “squarer” than in controls, indicating a 93-day longer healthspan (p = 0.02). PPP treatment was associated with toxicity (GI bleeding). Additional analysis of the drug likeness of NVP-ADW742 demonstrated potential cardiotoxicity and brain bioaccumulation. To conclude, small-molecule IGF1R inhibitors hold promise as a therapy that may improve human health span and lifespan; however, both molecules tested in this study have side effects that may outweigh their anti-aging effects.

**Statements and Declarations:** YB is an employee of ReGENE LLC. GB received compensation from ReGENE LLC as a consultant. CJE received compensation from and is a member of ReGENE LLC. AS received compensation from and is a member of ReGENE LLC. TS declares no conflict of interest.

## 1. Introduction

According to the antagonistic pleiotropy hypothesis of aging, if the same trait favors reproductive success at a young age but leads to the deterioration of health later in life, natural selection will support this trait due to a smaller contribution of older organisms to the next generation [1–3]. The most obvious situation when the same molecular mechanism has divergent effects at different ages of an organism occurs when the life cycle of the organism consists of distinct life stages [4]. For example, a mechanism increasing the fitness and survival of a caterpillar may have the opposite effect on the success of a butterfly. Humans, as well as other mammals, have two well-pronounced life stages in their postnatal life: the developmental life stage associated with reproductive immaturity and intensive growth, and the adult life stage associated with growth arrest and reproductive maturity. During the first stage, resources are mostly used for growth and development, while during the second stage, resources are used for energy accumulation in storage depots and reproduction [5]. Thus, the antagonistic pleiotropy hypothesis predicts that in mammals, aging mechanisms are heavily enriched with mechanisms of growth, development, and metabolic control.

This prediction is supported by extensive evidence from multiple animal models, which demonstrate that suppression of different nodes in the growth hormone/insulin-like growth factor/mechanistic target of rapamycin (GH/IGF/mTOR) cascade, the primary pathway regulating growth, development, and metabolism, increases longevity [6–11]. Additionally, many attempts to identify central mechanisms of aging in an unbiased way point to IGF signaling. For example, a manually curated database of aging genes, Open Genes, identifies 25 human genes linked to longevity and aging with high confidence [12]. More than half of these genes belong to the GH/IGF/mTOR cascade, and all major nodes of the cascade are part of this list. In another example, IGF-1 downregulation was identified as a central longevity mechanism within and across 41 mammalian species using multi-tissue transcriptomic data [13].

Antagonistic pleiotropy of IGF signaling is well recognized today and was reflected in the title of the Gordon Research Conference that took place in March 2025: *The Antagonistic Pleiotropy of IGF-1 and its Interactions with Insulin*. It is also supported by many lines of experimental evidence. For example, a clear biphasic effect of IGF1 receptor (IGF1R) on cardiac health was demonstrated, where mice with cardiomyocyte-specific overexpression of IGF1R exhibit enhanced cardiac function and exercise capacity at a young age but develop clear signs of heart failure at an older age; while young mice with reduced IGF1R signaling exhibit delayed cardiac growth but have attenuated cardiac aging [14, 15].

The coherent hypothesis linking the evolutionary and molecular roles of IGF signaling to aging makes this pathway an attractive target for anti-aging therapies. Indeed, IGF1R suppression by a monoclonal antibody was well tolerated in 18-month-old mice and resulted in significant improvements in health span and lifespan, including enhanced motor coordination and reduced systemic inflammation [16]. The life-extension effects of IGF-signaling suppression observed in model organisms have been corroborated by human studies. Population research has demonstrated that low IGF-1 levels predict greater life expectancy among exceptionally long-lived individuals [17, 18], while higher IGF-1 levels correlate with increased mortality and morbidity risks, including cognitive impairment [19]. Similarly, genetic mutations that attenuate IGF1R signaling [20] or reduce GH production were associated with extreme longevity [9]. Analysis of UK Biobank serum IGF-1 levels (*n* = 440,185) in relation to aging-associated diseases concluded an interaction consistent with antagonistic pleiotropy, where younger individuals with high IGF-1 are protected from disease, while older individuals with high IGF-1 face an elevated risk of developing disease or facing death [21].

Thus, the goal of this study is to test in a mouse longevity experiment if orally delivered small-molecule IGF1R inhibitors would be a promising candidate anti-aging therapy. We used two commercially available IGF1R inhibitors, picropodophyllin (PPP) and 5-[3-(phenylmethoxy)phenyl]-7-[trans-3-(1-pyrrolidinylmethyl)cyclobutyl]-7H-pyrrolo[3-d]pyrimidin-4-amine (NVP-ADW742). Both compounds, delivered via powdered diets, extended mouse healthspan, and NVP-ADW742 extended lifespan as well. Despite these positive effects, both molecules have poor drug-likeness properties, and new drug development efforts are needed to produce a new generation of small-molecule IGF1R therapies to be used as anti-aging therapies.

## 2. Materials and Methods

### 2.1. Diets and Treatment

Powdered PPP was purchased from Invivochem (cat. #: V062801) and NVP-ADW742 from Adooq Bioscience (cat. #: A10658). PPP is unstable at room temperature and transitions spontaneously to its highly toxic stereoisomer podophyllotoxin [22, 23]. Therefore, PPP was stabilized in accordance with the protocol described elsewhere [22] to reduce its degradation. In short, 3 g of PVP K30 (polyvinylpyrrolidone, Millipore Sigma, cat. #: 81420) were mixed with 8 mL of acetone (Millipore Sigma) and 2 mL of isopropanol (Fisher Scientific) until fully dissolved. 1 g of PPP was mixed with 18 mL of acetone until a homogenous slurry was obtained. Both mixes were combined and stirred vigorously for 30 min to form a suspension. The resulting suspension was evaporated overnight using a rotary evaporator at room temperature. Stabilized PPP was scraped from the flask and powdered using a mortar and pestle. NVP-ADW742 or stabilized PPP were mixed at required concentrations with irradiated powdered Teklad Global 14% protein rodent diet (Envigo, cat. # 2914CM) using Galaxy GMIX10 10 Qt. Planetary Stand Mixer (Main Street Equipment, cat. #: 177GMIX10). Stock diets were stored at 4°C.

In the pilot experiment, we used a range of doses of each compound, PPP and NVP-ADW742, to establish doses able to induce a systemic response in mice (see 2.2). To identify the range of doses for PPP, we used a reference oral dose of 3.2 mg of non-stabilized compound per mouse per day (approximately equal to 100 mg/kg body weight (BW)), a dose that was well tolerated for 16 days and produced significant systemic effects (positive effects on xenografts) in a previous study [24]. According to [22], the bioavailability of PPP increases 10-fold after stabilization. Therefore, we used 10 mg/kg BW of stabilized PPP as our highest dose. Four additional doses (1, 0.1, 0.01, and 0.001 mg/kg BW) were used to identify dose-response relations for systemic outcomes. There was only one study available by the start of our experiments, which reported oral administration of NVP-ADW742 in a mouse model [25]. This study demonstrated positive effects on xenographs at 50 mg/kg BW. Thus, 50 mg/kg BW was used as the highest dose of NVP-ADW742. Four additional doses (5, 0.5, 0.05, and 0.005 mg/kg BW) were used to identify dose-response relations for systemic effects. To achieve the desired dosing, we prepared diet mixes with the consideration that the average food consumption by adult C57BL6 mice is around 6g/30g BW per day [26].

### 2.2. Mouse Experiments

All animal experiments were conducted in the specific pathogen-free vivarium of the Baystate Research Facility. Mice were housed in ventilated polysulfone cages in an Allentown 140 cage rack with free access to water at controlled temperature (23 ± 2 °C), humidity (RH% 40 ± 10), and a 12 h dark/light cycle. Cages were provided with cotton nest pads (Nestlets, Ancare Corporation, Bellmore, NY, cat. #: NES3600) and ALPHA-dri® (Shepherd Specialty Papers, Portage, MI) bedding material free of endotoxins, mycotoxins, and environmental toxins, composed of virgin paper pulp. Mice were transferred to fresh cages every 14 days. All care and experimental procedures involving animals were approved by the Institutional Animal Care and Use Committee at the Baystate Research Facility (protocol #1727123).

#### 2.2.1. Pilot experiment

A pilot experiment was used to identify the optimal doses of compounds to be used in the longevity experiment. 55 male 70-day-old C57BL/6 (strain code 027) mice were purchased from Charles River Laboratories. After one week of acclimation, mice were randomly assigned to one of 11 dietary groups (n = 5/group): a control group and five different dose groups for each compound as described above. Mice were housed 5 per cage. Control and treatment powdered diets were provided ad libitum via four-ounce feeding jars with food followers and deluxe lids installed in cages (Dyets Inc, cat. # 910007, 910016, and 910024, respectively). Food followers were used to reduce diet loss from digging in jars. Feeding jars were inspected daily, and fresh diets were added if needed. Gnawing enrichment (Manzanita Wood Gnawing Sticks, cat.# W0016, BioServ) was provided in each cage. Following 7 days of treatment, all animals were euthanized, and samples of tissues expressing IGF1R at high levels (gonadal adipose, lung, kidney, and heart tissue) were collected and stored at -80°C. Blood serum was collected following 15 min centrifugation at 3000 g and stored at -80°C.

#### 2.2.2. Longevity experiment

75 male and 75 female 55-week-old C57BL/6 mice (stock #: 000664) were purchased from Jackson Laboratories. After one week of acclimation, mice were randomly assigned by use of a random number table to one of 6 dietary groups (n = 25/sex/treatment group) as follows: control male and female groups, 1 mg/kg BW of PPP male and female groups, and 0.5 mg/kg BW NVP-ADW742 male and female groups. Females were housed 3-4 per group, and males were initially housed 2 per group. Males were separated and single-housed in case of fighting, so that one month from the start of the experiment and till its end, approximately half of all males in each group were single-housed and half were housed in pairs. Treatment diets and enrichment materials were provided in the same way as in the pilot experiment. Throughout the whole experiment, animals were assessed daily for their body condition, activity, signs of abnormal behavior, injuries, and palpable anomalies, as described elsewhere [27]. All animals were weighed weekly. Severely moribund mice were euthanized for humane reasons. A mouse was considered severely moribund if it exhibited more than one of the following clinical signs following the NIH Interventions Testing Program protocols: (i) severe lethargy, as indicated by reluctance to move when gently prodded; (ii) inability to eat or to drink; (iii) rapid weight loss over 1 week or more; or (iv) a severely ulcerated or bleeding tumor [28]. The age at which a moribund mouse was euthanized was taken as the best available estimate of its natural lifespan. Mice found dead were also noted at each daily inspection. The experiment was terminated when mice reached 151 weeks of age. Thirteen animals out of the 150 survived to this age. Each mouse found dead or euthanized was subjected to gross necropsy to identify the plausible causes of death or moribund condition. Throughout the longevity experiment, mice were subjected to a range of tests and measurements (see below), which were performed between 8 and 11 am.

#### 2.2.3. Heart rate and blood pressure assessment

Heart rate and blood pressure were measured in 62-week-old mice using the BP-2000 Blood Pressure Analysis System (Visitech, BP-2000-M-6) as recommended by the manufacturer. This tail-cuff blood pressure measurement is a non-invasive test using the same principle as that of the inflatable blood pressure cuff used in humans. Mice were trained for two consecutive days, with the number of measurement cycles set at 5 for each training session. This training allowed mice to habituate to the device and procedure and reduce the noise signal. The testing session was performed on day three and included 10 measurement cycles, from which the final readings were averaged.

#### 2.2.4. Assessment of short-term memory

Short-term memory was assessed in 65 and 98-week-old mice using Y-maze spontaneous alternation protocol as described in detail elsewhere [29]. This test was validated for the pharmacological investigation of memory [30]. It is based on natural exploratory behavior, which favors entrances to “unexplored” arms of the maze rather than reentering the same or recently visited arm. Deviation from sequential entries to all three arms is considered a “mistake” of short-term memory. For this task, mice were placed into the designated start arm facing the center of the maze with no visual cues. The subject was evaluated for alternation behavior during an 8-minute observation session, and its behavior was recorded and tracked to provide the number and sequence of arm entries. The Y-maze is a plastic arena with the following dimensions of each arm: 33.65 cm length, 6 cm width, and 15 cm height of walls.

#### 2.2.5. Kidney function assessment

Kidney function was assessed by the urine ratio of microalbumin to creatinine [31] in 68-week-old mice. Urine was collected in the morning hours for three consecutive days, and pooled samples were analyzed using the Albumin-to-Creatinine Ratio (ACR) Assay Kit (BioVision, cat. #: K551).

#### 2.2.6. Grip strength assessment

Grip strength was analyzed in 70- and 131-week-old mice using a Grip Strength Meter (Ugo Basil, Cat. #: 47200) to assess neuromuscular function as described before [29]. The task was performed by allowing a mouse to grab onto a wire mesh grid with all four paws, then pulling the mouse gently but firmly by the tail until it released its grip from the grid. A force transducer was measuring the maximum force (measured in grams). In one round of grip strength assessment, each mouse was tested for three consecutive trials with 20-second intervals, and the highest measured force was recorded.

#### 2.2.7. Assessment of motor coordination

Motor coordination of 72-week-old mice was assessed using the Rotarod assay following the protocol for aged mice [29]. The Rotarod (Ugo Basile) consists of a computer-controlled, motorized horizontal rotating rod. Each mouse was first habituated to the device and procedure via two consecutive 2-minute training sessions with the rod rotating at a constant speed of 4 revolutions per minute (rpm). In a testing session, the subject was placed on a rod that began rotating at four revolutions per minute (rpm) and accelerated over 5 min to a maximum of 40 rpm. The latency to fall was recorded as an indicator of motor coordination. All intervals between training and testing sessions were 90 min.

#### 2.2.8. Assessment of hair graying

At the start of the longevity experiment, all mice had black shiny fur; however, after approximately three months of treatment, the team of the project noticed a potential effect of treatments on hair graying. The following protocol was developed to quantify hair graying, and quantification was done in 73-week-old mice. An image of fur behind the base of the right forelimb was taken using a digital camera (see Fig. 1S in the Supplementary Information). Images were analyzed on a computer using Fiji (Image J) software as follows. A field of 1 cm sq was selected and converted to a 16-bit shades of gray image. Next, the MaxEntropy threshold was set to convert it to black and white. After inversion, the percentage of area occupied by black pixels was measured. This percentage represented the area occupied by gray hair.

#### 2.2.9. Depression-related phenotype assessment

79-week-old mice were subjected to the tail suspension test to assess depression-related phenotype. Tail suspension places mice in a temporary inescapable stress situation, which is interpreted as learned helplessness or behavioral despair [32, 33]. The mouse initially attempts to escape but eventually ceases in an immobility response. The longer duration of initial attempts to escape indicates a more resilient phenotype. An apparatus consists of a stand with a horizontally mounted bar, 30 cm above the table surface. The test was performed as described in detail elsewhere [31]. Individual mice were suspended by a point within the last 2–3 cm from the tip of the tail on the stand bar and secured using masking tape. Mice were monitored for five minutes, or till the moment when they stopped attempting to escape for 10 consecutive seconds, whichever happened earlier.

#### 2.2.10. Glucose tolerance test

We assessed how rapidly glucose is cleared from the circulation in 80-week-old mice. Test parameters were selected based on the study, which evaluated the efficiency of the test in relation to different modifications of the protocol [34]. Mice were fasted for 6 hours before the test, and 20% glucose solution in a sterile 0.9% saline was injected intraperitoneally using a 25G 5/8 needle in an amount equivalent to 2 g glucose per kg body weight. Blood glucose was measured in the tail vein using the Contour Next ONE blood glucose monitoring system (Ascensia Diabetes Care) before glucose injection and 30, 60, and 120 min after.

#### 2.2.11. Frailty assessment

Frailty assessment was performed in 89-week-old mice following the protocol described in detail elsewhere [29]. In short, mice were individually observed and evaluated for the absence or presence and severity of 27 different characteristics, each scored as 0, 0.5, or 1 based on the level of severity: signs of alopecia excluding barbering around the snout, loss of fur color, dermatitis or presence of skin lesions, whiskers density, coat condition, signs of piloerection, defined as erection of the hairs, eyes for cataracts indicated by opaque spots, eyes for discharge or swelling, size of the eyes for microphthalmia, eyes for cataracts as indicated by clouding, nares and snout for nasal discharge, rectal area for prolapse, penile (male) or vaginal (female) prolapse, rectal area for diarrhea, body for the presence of tumors, kyphosis as indicated by hunched posture due to curvature of the spine, sacroiliac region (back and pubic bones) to evaluate body condition, gait, sign of tremor, breathing rate and depth, menace reflex, tail stiffening, vestibular disturbance, extension of the forelimbs for visual placing, distended abdomen, teeth for uneven position or malocclusion, and righting reflex response.

### 2.3. Molecular markers of systemic IGF1R inhibition

Free blood IGF-1 was measured in murine serum using RayBio® Mouse IGF-1 ELISA Kit (Catalog #: ELM-IGF1). Serum samples were diluted 100-fold and measured in accordance with the manufacturing guidelines. Total RNA was isolated from gonadal adipose, lung, kidney, and heart tissue using TRIzol reagent (Invitrogen) and was quantified using a NanoDrop 1000 instrument (Thermo Fisher Scientific, Wilmington, DE). Total RNA was reverse transcribed using the iScript™ Reverse Transcription Supermix for RT-qPCR (Cat. # 170-8841, BioRad). Triplicate 5-μl RT-qPCR reactions, each containing iTaq Universal SYBR Green Supermix (Cat. # 172–5124, BioRad), primers, and cDNA template, were loaded onto a 384-well plate and run through 40 cycles on a CFX384 real-time cycler (Bio-Rad Laboratories, Inc). The data were analyzed using the manufacturer’s CFX manager software, version 3.1. Relative quantification was determined using the ΔCq method and presented in % change in comparison with the control group, where the level of expression in the control group is 100%. RT-qPCR was conducted with primers for genes regulated downstream of IGF-1 or members of the IGF cascade: Socs3, Osgin1, Cxcl, Hspa8, Rgs5, Igfbp3, Igf1, Serpina, and Vegfa [35–39]. B2M was used as a housekeeping gene due to its validated expression stability across tissues [40, 41]. Primers were designed using the UCSC genome browser and Primer3Plus software so that forward and reverse primers anneal to different exons spanning a long intron (Table 1S in the Supplementary Information).

### 2.4. Drug-likeness properties of NVP-ADW742

Because NVP-ADW742 demonstrated improved mouse health span and lifespan in our longevity experiments, we characterized its drug-likeness properties to assess whether it has a favorable profile for clinical application. Specifically, NVP-ADW742’s physical-chemical properties were calculated using the ACDlabs software (Advanced Chemistry Development, Inc., Toronto, Ontario, Canada). Additionally, water solubility, IGF1R dissociation constant (Kd), and half-maximal inhibitory concentration (IC50) for IGF1R and insulin receptor (IR) were determined using services provided by Eurofins (Eurofins Discovery Services North America, LLC, St Charles, MO, and Eurofins Panlabs Discovery Services Taiwan Ltd). In-vitro and in-vivo pharmacokinetic studies, as well as in-vitro toxicity using HepG2 cell viability assay and hERG assay, were performed by WuXi AppTec Ltd (Kowloon, Hong Kong). Methods used by contract research organizations are described in Supplementary Methods.

### 2.5. Statistical methods

Each mouse originally entered the study was, at the time of analysis, considered to be in one of two categories: either dead (from natural causes) or censored. Mice were censored at the age when they were no longer subjected to the mortality risks, including mice that died as a result of an accident, and 13 mice that were euthanized at the date of the experiment termination. Kaplan– Meier analysis considered censored mice to be lost from follow-up on the day on which they were removed from the longevity protocol. Unless stated otherwise, all significance tests about survival effects are based upon the two-tailed log-rank (Mantel-Cox) test at *p* < 0.05, with censored mice included up until their date of removal from the longevity population. Statistical claims related to maximum lifespan are based on the procedure of Wang *et al*. [42], which uses Fisher’s exact test to compare the proportions of surviving mice, in control and test groups, at the age corresponding to the 90th percentile for survival in the joint distribution of the control and test groups together. A similar approach was used to identify differences in healthspan, where Fisher’s exact test was used to compare the proportions of surviving mice in control and test groups, at the age corresponding to the 20th percentile for survival in the joint distribution. All data for behavioral and physiological tests are shown as mean ± SE, and the *p*-value is for the T-test comparison with controls of the corresponding sex without adjustment for multiple comparisons.

## 3. Results

### 3.1. Identification of treatment doses in a pilot study

First, we identified which molecular markers may be used as indicators of the systemic effects of IGF1R inhibitors. The most pronounced response to PPP and NVP-ADW742 treatment was found for Vegfa expression in gonadal adipose tissue and free IGF-1 in blood serum. These markers were further used to develop dose-response curves – see Fig. 1. Dose-response curves were U-shaped for Vegfa expression for both IGF1R inhibitors, and expression was significantly different from controls in a range of 0.05-50 mg/kg BW for NVP-ADW742 and in a range of 0.01-1 mg/kg BW for PPP. Free serum IGF-1 decreased significantly at all tested doses of NVP-ADW742, higher than or equal to 0.05 mg/kg BW, and at doses of PPP higher than or equal to 1 mg/kg BW. Using these results, we selected for the longevity experiment the minimal daily doses that reached the highest response of both markers of IGF1R inhibition. These doses are 0.5 mg/kg BW/day of NVP-ADW742 and 1 mg/kg BW/day of PPP.

**Fig. 1.**
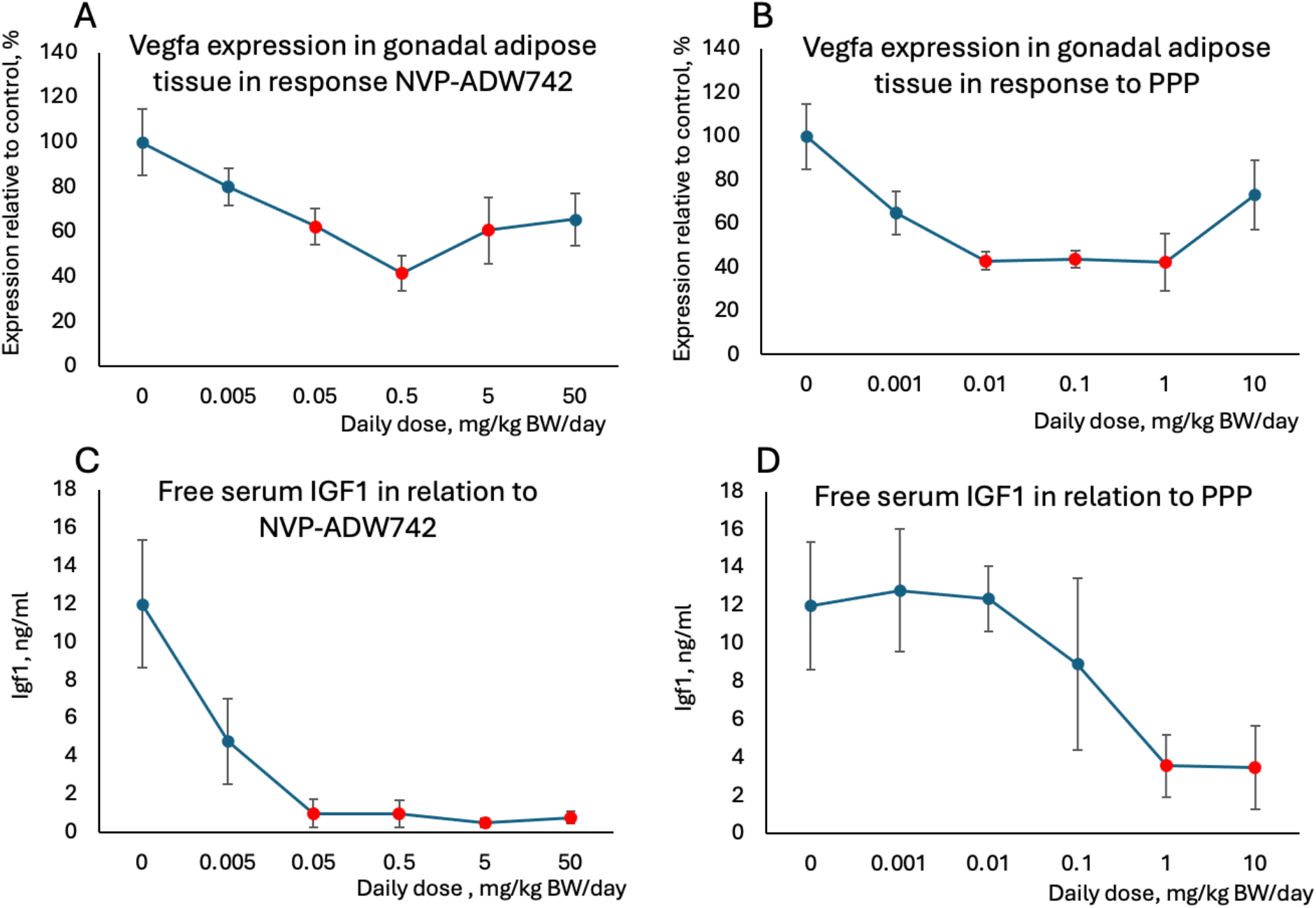
Dose-response curves for Vegfa expression in adipose tissue (A and B) and free serum Igf-1 (C and D) in response to dietary exposure to different doses of NVP-ADW742 (A and C) and PPP (B and D). Each datapoint on the graphs is mean ± SE (n=5/group). Red datapoints are significantly different from controls (T-test p ≤ 0.05).

### 3.2. IGF1R inhibition improves cognitive and neuromuscular functions

In humans, aging is often associated with progressive loss of short-term memory, development of depression, loss of neuromuscular coordination, and physical strength. To test if these health parameters are affected by IGF1R inhibition, we conducted the following tests, respectively: spontaneous alteration, tail suspension, rotarod, and grip strength.

Short-term memory assessment was done twice: at 9 and 41 weeks after the start of treatment (Fig. 2 A,B). At the first time point, short-term memory demonstrated improvement in both males and females treated by both compounds, as compared with the same age control animals, although this shift was not significant for most groups (p = 0.08 and 0.06 in females, and p = 0.008 and 0.06 in males treated with NVP-ADW742 and PPP, respectively). When short-term memory was remeasured 41 weeks after the start of treatment (> 2-year-old mice), a significant decline was observed in both male and female control mice, while both treated groups were mostly protected from this decline. The difference between treated and control mice reached significance for most groups (p = 0.003 and 0.03 in females, and p = 0.02 and 0.06 in males treated with NVP-ADW742 and PPP, respectively). During the short-term memory assessment at 65 weeks of age, around 17% of animals attempted to mount the maze wall during the test. This number dropped to 7% by 98 weeks of age. Both male and female NVP-ADW742-treated animals were more active in wall mounting at both time points than the control and PPP groups (Table 2S in the Supplementary Information). This difference was significant in 65-week-old mice (*X*^2^(2, N=146) = 5.7, *p* = 0.05).

**Fig. 2.**
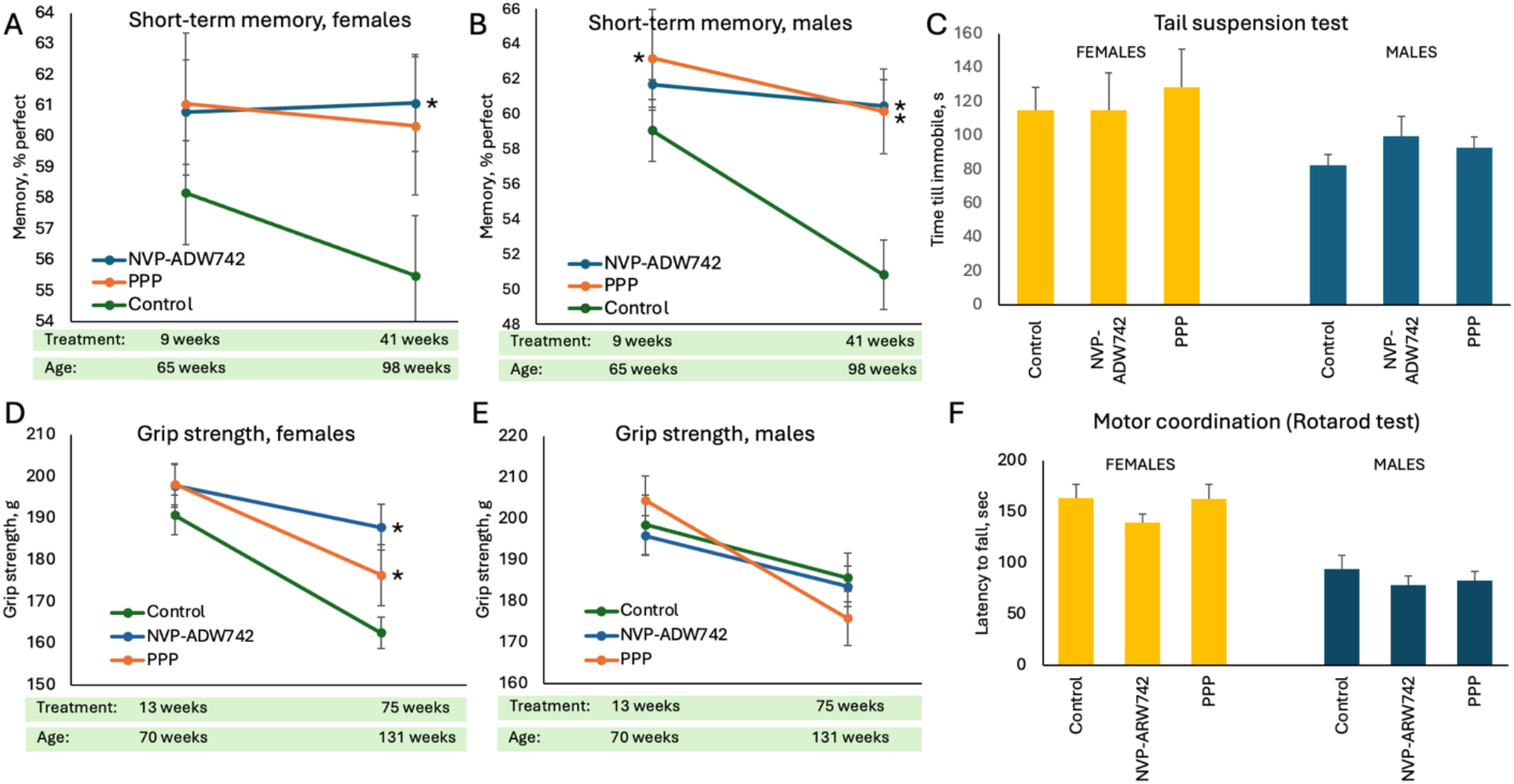
Effects of long-term treatment with IGF1R inhibitors on behavioral and neuromuscular functions. A and B – short-term memory assessed in 65- and 98-week-old female and male mice, respectively, using the spontaneous alterations test. C – assessment of depression-like phenotype using the tail suspension test in 79-week-old mice. D and E – grip strength of 70- and 131-week-old female and male mice, respectively. F – motor coordination assessed in 72-week-old mice using the Rotarod test. All data are mean ± SE, stars indicate p-value < 0.05 for T-test comparison with the same age and sex controls.

There were no significant differences between treatment groups and controls in the time 79-week-old mice spent trying to escape before reaching an immobile state in the tail suspension group (Fig. 2C). Overall, across all groups, females demonstrated less “depressive” phenotype (p = 0.02) than males, with 91.5 ± 8.2 and 119.5 ± 19.1 sec before immobility in males and females respectively (mean ± SE). Grip strength was declining with age in all groups of mice (Fig. 2 D,E); however, IGF1R inhibitor treatment demonstrated some protective effect for this decline in females. This trend was already seen after thirteen weeks of treatment, but it reached significance only after 75 weeks of treatment, in 2.5-year-old mice (p = 0.005 and 0.05 for NVP-ADW742- and PPP-treated females, respectively). No significant differences were found between treatment groups in the Rotarod test (Fig. 2F).

### 3.3. IGF1R inhibition affects metabolic and cardiovascular health

In the longevity experiment, changes in body weight followed similar patterns across all treatment and sex groups. This included an initial drop during the adaptation period, a gradual increase until around 20 months of age, a plateau until about 25 months of age, and a subsequent decline (Fig. 3A). Males in both treatment groups gained weight more significantly during the increase phase and experienced a faster weight loss during the decline phase. In contrast, females in the treated groups maintained higher body weight than those in the control group for most of the experiment. No differences in glucose tolerance were seen in response to IGF1R inhibitors in 80-week-old female mice after 24 weeks of treatment. All groups of females exhibited a glucose response similar to that of young healthy animals (Fig. 3B), with blood glucose reaching levels close to fasting values at 120 minutes [43]. Conversely, control males showed a diminished response to glucose (Fig. 3C). These findings align with the established age-dependent reduction in glucose tolerance seen in male but not female C57BL/6 mice [44]. Importantly, both IGF1R inhibitors partially mitigated this decline. Measurements of blood pressure and cardiac pulse in 62-week-old mice, following six weeks of treatment, showed a decrease in systolic blood pressure in males (Fig. 3E), but not in females (Fig. 3D), along with a reduction in pulse beats per minute in both sexes for both groups receiving IGF1R inhibitors (Fig. 3F). To assess renal function, we measured the albumin/creatinine ratio in 68-week-old mice, finding no differences between the control and treated groups. Across all groups, the values for albumin, creatinine, and their ratio were 7.95±0.65 mg/dL, 0.047±0.002 mg/dL, and 225.7±29.7 in male mice, and 9.17±0.68 mg/dL, 0.058±0.002 mg/dL, and 169.0±13.2 in female mice, respectively.

**Fig. 3.**
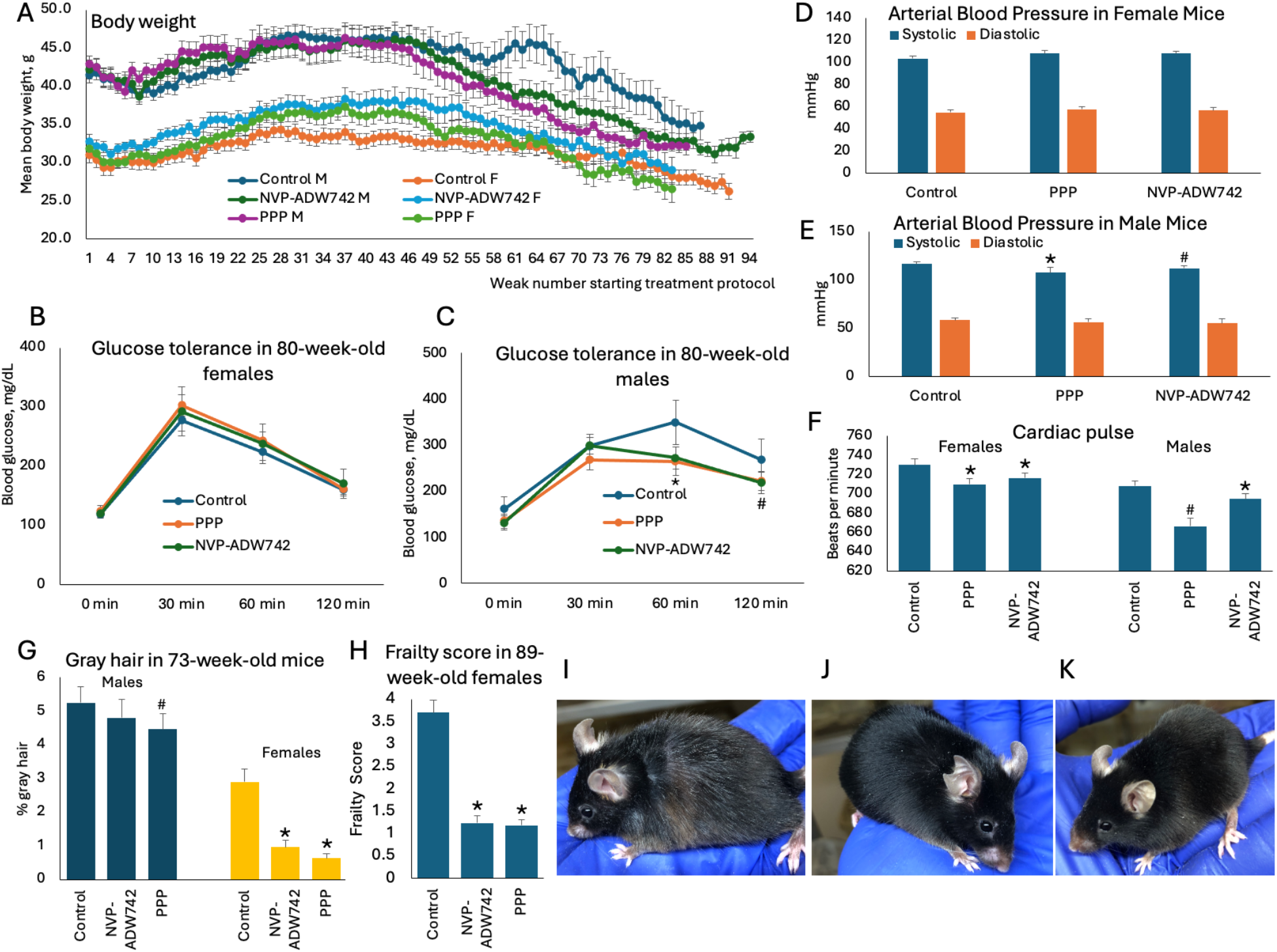
Effects of long-term treatment with IGF1R inhibitors on aging phenotype. A -body weight changes across the treatment period, starting at 56 weeks of age. Data are shown for every group until their sample size dropped below 5 animals. B and C – glucose tolerance test in 80-week-old female and male mice, respectively. D and E – Arterial blood pressure in 62-week-old female and male mice, respectively. F – cardiac pulse in 62-week-old mice. G – percent gray hair in 73-week-old mice. H – frailty score in 89-week-old females. I, J, and K - Representative images of control, NVP-ADW742, and PPP-treated 73-week-old females, respectively. In graphs, all data points are shown as mean ± SE; * p < 0.05, ^#^ p < 0.09.

### 3.4. IGF1R inhibition attenuates the aged phenotype development

At the start of the longevity experiment, all mice had black, shiny fur; however, after approximately three months of the treatment protocol, some mice began to develop gray hair. To assess the degree of hair graying, we quantified the area occupied by gray hair in fur images in 73-week-old mice. Treatment reduced hair graying non-significantly in male mice (Fig. 3G) while both treatments significantly reduced hair graying in females (Fig. 3G,I,J,K and Fig. 2S) with percent fur occupied by gray hair 2.9±0.4 in controls, 0.9±0.2 (*p* = 4.7E-05) in NVP-ADW742 group, and 0.6±0.1 (p = 1.2E-06) in PPP treated group. Although not quantified at later time points, hair graying occurred in all control females, while females in both treated groups maintained black and shiny fur till death. Frailty score based on 27 phenotypic parameters was assessed in 89-week-old mice. No difference in frailty scores was observed among the treatment groups of male mice. Among female mice, frailty scores were around 3 times higher in the control group (3.7±0.3) as compared with NVP-ADW742 (1.2±0.2, p=1.8E-09) and PPP (1.2±0.1, p=6.5E-10) groups (Fig. 3H). Higher scores in control females were dominated by a higher incidence of poor skin and fur conditions (gray hair, ruffled and ungroomed fur, alopecia, dermatitis, and loss of whiskers), rectal prolapse, vision loss, and menace reflex.

### 3.5. Survival analysis and inferred causes of death

Survival curves within each treatment group had a similar appearance for male and female mice and were not statistically different (Fig. 4A-C). Therefore, we combined data from both sexes and made all comparisons between treatment groups for these combined groups (Fig. 4D). No significant differences between groups were observed in Kaplan–Meier analysis. However, mean longevity values were higher in NVP-ADW742-treated mice (892.3 days, SD = 131.9, T-test p = 0.06) than in controls (848.8 days, SD = 152.0) with a 43.5-day mean difference. Although non-significant, mean longevity values were lower in PPP-treated mice than in controls (824.4.3 days, SD = 176.4, T-test p = 0.46). The difference between survival curves in the control group and the NVP-ADW742 group was mostly due to a “squarer” survival curve (longer health-span) in the NVP-ADW742 group than in the control mice. To test this difference statistically, we used a modification of the procedure for maximum lifespan comparison [42]: Fisher’s exact test was used to compare differences between treatment groups in the numbers of mice with lifespans above and below the 20^th^ percentile. NVP-ADW742-treated mice had 93 days longer healthspan than controls (p = 0.02), indicating compression of the morbidity period and the shift in the average lifespan towards the maximum lifespan. No difference was observed between healthspan in the control and PPP groups. Similarly, no differences in the maximum lifespan were observed between experimental groups. Gross necropsy provided plausible causes of death or severe moribund conditions that resulted in euthanasia (Table 3S in the Supplementary Information). The leading lesions present in >5% of animals were as follows: neoplasia – 65.8%, including tumors of liver (26.8%), lungs (12.1%), pancreas (8.1%), spleen (7.4%), and brain (5.4%); abscess and/or necrosis of seminal vesicles – 31.1% of male mice; liver failure presumed when liver was 3-5 times enlarged, contained nodules and/or tumors, cysts, and overfilled gull bladder often containing abnormally colored liquid – 30.2%; heart failure as presumed by the presence of pleural effusion and/or abnormal heart appearance – 16.1%; urinary obstruction seen as overfilled bladder and visible bladder stones – 13.4%; and gastrointestinal bleeding – 5.4%. The patterns of pathology were very similar between treatment groups, with the following exceptions: NVP-ADW742-treated mice had a significantly lower (*X*^2^ (2, *N* = 149) = 8.57, *p* = 0.01) incidence of urinary obstructions (1 case) than control group (9 cases) and PPP-treated mice (10 cases); and PPP-treated mice had a significantly higher (*X*^2^ (2, *N* = 149) = 6.79, *p* = 0.03) incidence of gastrointestinal bleeding (6 cases) than other groups (1 case in each). Most cases of gastrointestinal bleeding in the PPP group occurred early, and therefore, they contributed to a flatter shape of the curve than in controls at the beginning of the mortality period.

**Fig. 4.**
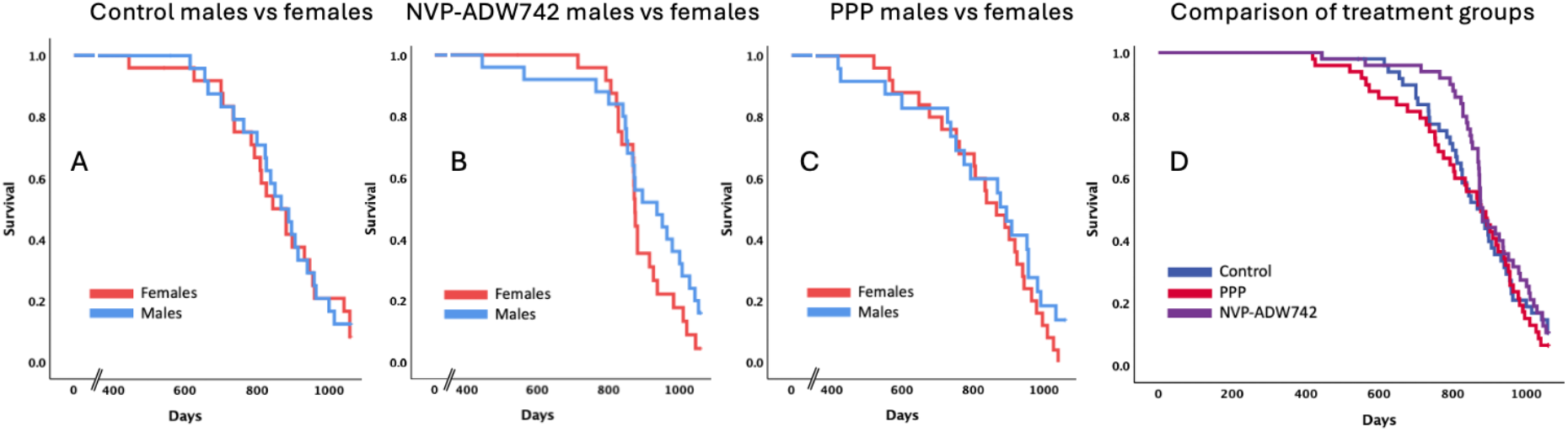
Survival curves in the control group (A) and groups treated with NVP-ADW742 (B) or PPP (C). D – survival curves for treatment groups combined for male and female mice.

### 3.6. Drug-likeness properties of NVP-ADW742

NVP-ADW742 properties are summarized in Table 1. Some of these properties indicate an unfavorable profile for clinical application as an oral drug. Specifically, NVP-ADW742 has high affinity in the hERG assay (IC_50_ < 300 nM). The hERG assay (human ether-a-go-go-related gene assay) evaluates a drug’s effects on the electrical abnormalities in hERG potassium channel function, a known risk factor for arrhythmias and cardiac safety [45]. Additionally, the drug has low bioavailability (F = 4.5%) and rapid unbound plasma clearance as indicated in Table 1. The low bioavailability correlates with low Caco-2 permeability and high efflux. More data is needed, but the longer T_1/2_ for human versus mouse microsomes suggests an improved human in vivo PK profile. Nonetheless, efforts to increase drug exposure will need to include understanding and mitigating clearance. Finally, NVP-ADW742 sequesters in the brain as indicated by higher brain than plasma AUC and protein binding. This property likely ensured a positive effect of NVP-ADW742 on brain function in our experiments, despite its overall low bioavailability. However, brain bioaccumulation may make control of brain dose highly challenging.

**Table 1.** NVP-ADW742 drug-likeness properties. Caco-2 P_app_ A-B - apparent permeability coefficient in apical-to-basolateral transport across a confluent monolayer of Caco-2 cells; Cl_u_ – unbound drug clearance, F – oral bioavailability, fAUC – total free drug area under the curve, fC_max_ – maximum free drug concentration, IC50 – half-maximal inhibitory concentration, IV – intravenous, Kd – dissociation constant, K_p,uu,brain_ – unbound brain to plasma partitioning coefficient, LogD – computed distribution coefficient for compound ionized at pH 7.4, LogP – computed distribution coefficient for non-ionized compound, MW – molecular weight, pKa – computed acid dissociation constant, PO – peroral, PPB – plasma protein binding, T_1/2_ – half-life, TPSA – computed topological polar surface area, V_ss_ – steady state volume of distribution.

| Target Affinities |  |
| --- | --- |
| IGF1R Kd (nM) | 8.6 |
| IR IC50 (nM) | 350 |
| IGF1R IC50 (nM) | 39 |
| Toxicity |  |
| HepG2 IC50 (nM) | 9000 |
| hERG IC50 (nM) | <300 |
| Human ADME |  |
| Human microsomes $T_{1/2}$ (min) | 67.8 |
| Caco-2 $P_{app}$ A-B ($10^{-6}$ cm/s) | 0.45 |
| Caco-2 efflux ratio | 11.6 |
| Mouse DMPK Properties |  |
| Mouse microsomes $T_{1/2}$ (min) | 9.9 |
| IV $Cl_u$ (mL/min/kg) | 5700 |
| IV $V_{ss}$ (L/kg) | 6.8 |
| PO $fC_{max}$ (ng/mL) | 0.35 |
| PO $T_{1/2}$ (h) | 1.9 |
| PO fAUC (ng*h/mL) | 1.23 |
| Oral F (%) | 4.5 |
| Plasma % free | 1.4 |
| Mouse Plasma and Brain PK |  |
| Plasma AUC ng*mL/h | 331 |
| Brain AUC ng/g/h | 461 |
| PPB % | 98.6 |
| Brain homogenate binding % | 99.8 |
| Plasma fAUC (ng*h/g) | 4.6 |
| Brain fAUC (ng*h/g) | 0.9 |
| $K_{p,uu,brain}$ | 0.2 |
| Physiochemical Properties |  |
| MW | 453.6 |
| Solubility water (μM) | 132 |
| LogP (calc) | 4.6 |
| LogD (calc) | 2.1 |
| pKa (calc) | 10 |
| TPSA | 69 |

## 4. Discussion and conclusions

This study demonstrates that two small-molecule IGF1R inhibitors (PPP and NVP-ADW742) delivered with food starting at 13 months of age improved several indices of healthspan in aging mice, and NVP-ADW742 altered the shape of the survival curve, indicating compression of the aging-associated morbidity period and the shift of an average lifespan towards the maximum lifespan. Specifically, both compounds protected both sexes from short-term memory decline; reduced systolic blood pressure in males and pulse rate in both sexes; rescued declining glucose tolerance in males; and abolished grey hair development, reduced frailty, and protected against grip strength decline in female mice. Importantly, we observed improvement in many healthspan indices in both sexes. These findings indicate that small-molecule IGF1R inhibitors hold promise as a therapy that may improve human health span.

The involvement of genes homologous to the mammalian IGF1R (InR in *Drosophila melanogaster* and DAF-2 in *Caenorhabditis elegans*) in lifespan regulation was first discovered in invertebrates [46–48]. Initial attempts to modulate mammalian longevity by IGF1R suppression used heterozygous knockout mice for Igf1r and produced conflicting results. For example, in one study, heterozygous mice lived on average 26% (33% in female and 16% in male mice) longer than their wild-type littermates and did not develop any adverse phenotype [49], while another study reported only a slight increase (< 5%) in mean lifespan of females and no increase in males [50]. Additionally, in the later study, aged mice developed insulin resistance, and males also developed glucose intolerance. Today, antagonistic pleiotropy of IGF signaling is well recognized [21, 51– 53], suggesting that the use of heterozygous Igf1r knockouts in aging research may have low relevance, and manipulations targeting IGF signaling only in late adulthood are a preferred approach. Indeed, IGF1R suppression by a monoclonal antibody starting at 18 months of age improved several indices of healthspan, reduced death from neoplastic disease, and increased mean and median lifespan in females [16].

Historically, clinical interest in IGF1R inhibitors was mostly motivated by the development of anti-cancer therapies. Multiple IGF1R antibodies were explored (e.g., CP-751,871, AVE1642/EM164, IMC-A12, SCH-717454, BIIB022, AMG 479, MK-0646/h7C10) [54], including clinical evaluation of anti-IGF1R drugs for cancer treatment. These studies showed no significant benefit, and most pharmaceutical companies have abandoned their IGF1R drug development programs [55]. Anti-IGF1R antibody teprotumumab was approved by the US Food and Drug Administration in 2020 to treat thyroid eye disease [56]. Antibody-based therapies have unfavorable profiles as potential anti-aging prophylactic therapies due to inherent challenges associated with biologics delivery in general and due to their poor ability to cross the blood-brain barrier in particular [57–59], as well as the long half-life of antibodies in circulation, which may be associated with adverse side effects [54, 60, 61]. Therefore, in the current study, we used commercially available small-molecule inhibitors of IGF1R: PPP and NVP-ADW742.

Neither compound is optimized for clinical use, and their adverse side effects may likely outweigh the anti-aging effects of IGF1R inhibition. Specifically, PPP is a cyclolignan unstable at room temperature that transitions spontaneously to its highly toxic stereoisomer podophyllotoxin [22, 23]. Podophyllotoxin is lipid soluble, so it crosses the cell membrane easily and inhibits mitotic spindle formation, resulting in neurotoxicity, bone marrow depression, gastrointestinal irritation, and hepatic and renal dysfunction [62]. Podophyllotoxin contamination is a plausible explanation for the increased gastrointestinal bleeding cases in the PPP group as compared with the control and NVP-ADW742 groups. It is also possible that the decreased urinary tract obstruction incidence seen in the NVP-ADW742 group, as compared with controls, was not seen in the PPP group due to the renal dysfunction induced by podophyllotoxin. Similarly, reduced average lifespan in the PPP group may be due to podophyllotoxin toxicity. This assumption is supported by the fact that most deaths at the beginning of the mortality period in the PPP group were due to gastrointestinal bleeding, a cause that had a negligible role in other groups.

Because NVP-ADW742 demonstrated improvement of mouse healthspan and lifespan, we characterized this compound for its drug-likeness properties. However, the drug-likeness profile of the compound is also not favorable for clinical application. Specifically, NVP-ADW742 suppresses the hERG potassium channels at low doses, indicating potential arrhythmias and cardiac toxicity, has low bioavailability and rapid clearance, and bioaccumulates in the brain, making control of brain dose highly challenging. NVP-ADW742-treated mice had a higher, although non significantly, incidence of heart failure concordant with in-vitro hERG binding. Additionally, cardiotoxicity may be difficult to register at gross necropsy, resulting in an underestimate. Thus, we conclude that although small-molecule selective IGF1R inhibitors bear promise as longevity therapy, both compounds tested in the current study have major flaws, making them unfavorable candidates for clinical testing.

This study has several limitations. For example, higher n per treatment/sex group may identify additional differences in survival and plausible causes of death. Additionally, aging is associated with changes in almost every function and numerous molecular markers, while only a few were assessed in this study. Only single drug doses were investigated, limiting the understanding of dose-response relationships. Finally, the exact molecular mechanisms linking IGF1R modulation with the aging process remain poorly understood.

