## Supplementary Information for "Small-molecule IGF1R inhibitors extend healthspan in a mouse model"

### GeroScience

### SUPPLEMENTARY FIGURES

**Figure 1S.** Fiji (Image J) pipeline for quantification of grey hair percentage

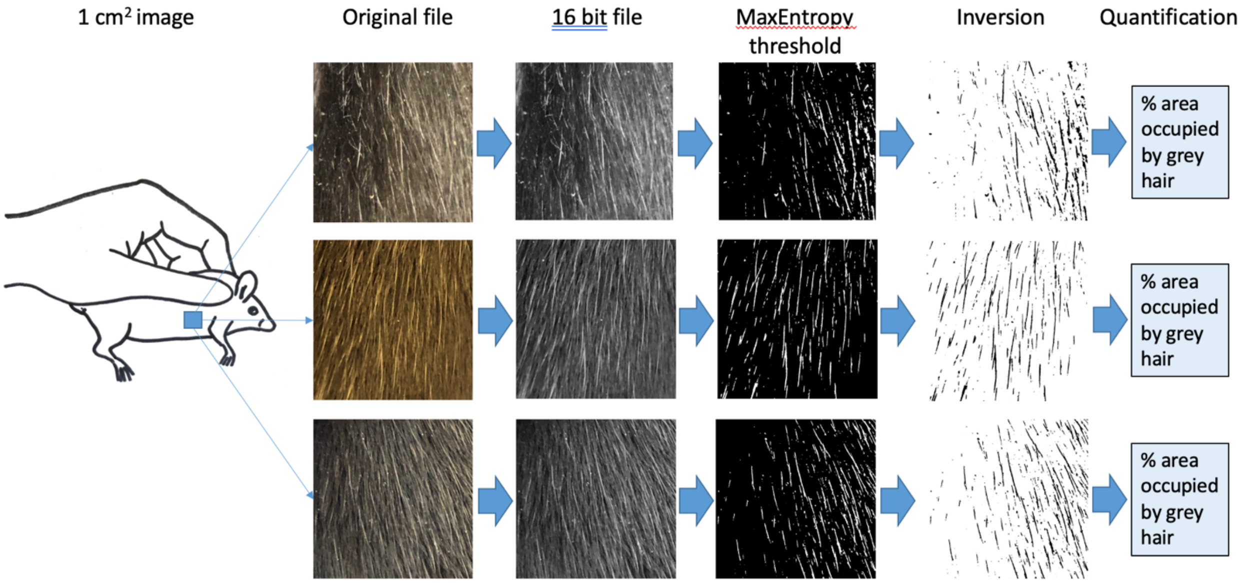

**Figure 2S.** Representative images of 73-week-old female C57BL/6J mice following 17 weeks of treatment.

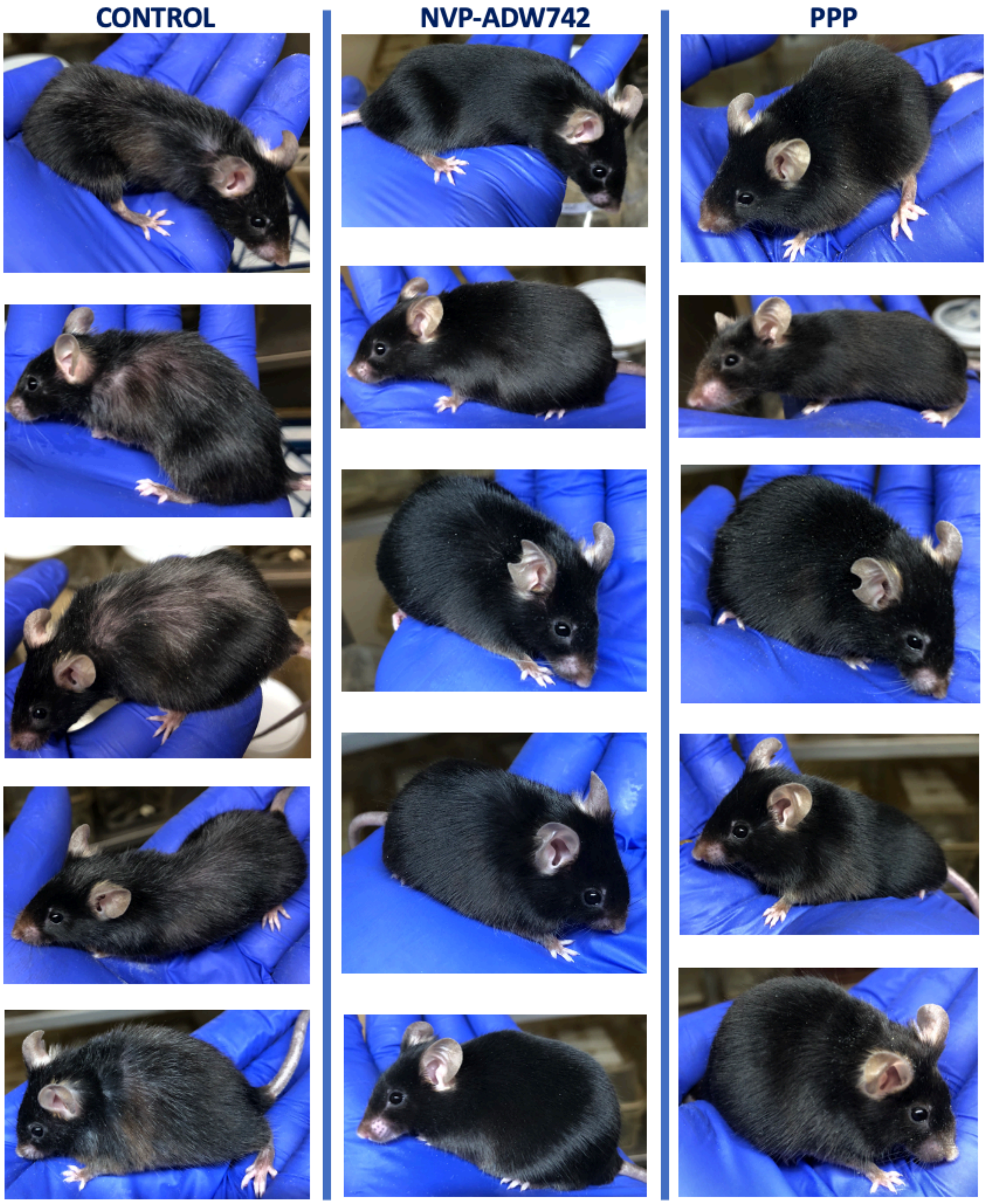

**Figure 3S.** Voltage command protocol for hERG channel test on SyncroPatch

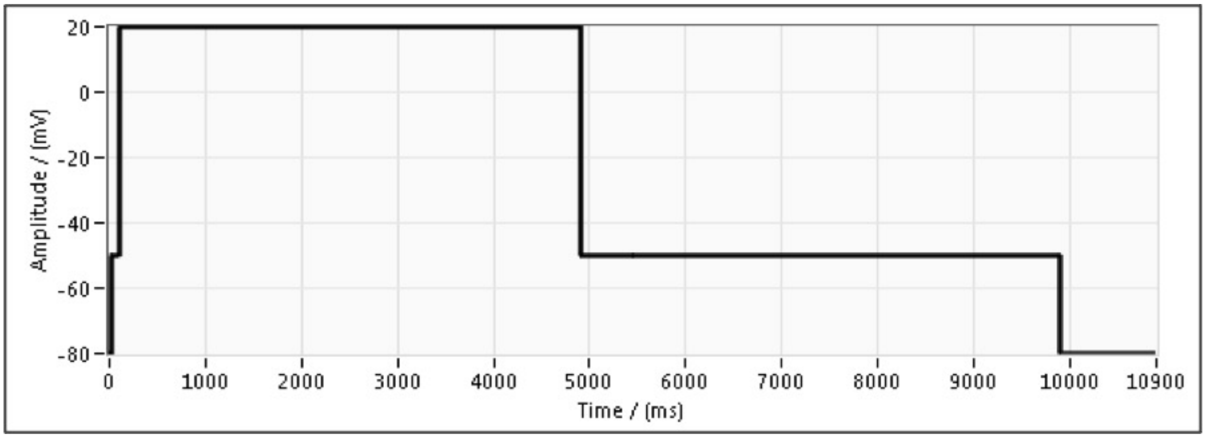


### SUPPLEMENTARY TABLES

**Table 1S.** Description of qRT-PCR primers

| Gene | Primers (5'-3' sequence) | Exon | Amplicon | Comments |
|---|---|---|---|---|
| B2m | CCGGCCTGTATGCTATCCAG<br>TGTTCGGCTTCCCATTCTCC | 1<br>2 | 78 | Housekeeping gene |
| Socs3 | GAGATTTCGCTTCGGGACTA<br>AACTTGCTGTGGGTGACCAT | 1<br>2 | 129 | Gene is responsive to IGF-1 [1–3] |
| Osgin1 | ACCAGCTCCCTCCAGCTATC<br>ACTACTGGGAGGGGCTCTGA | 1<br>2 | 133 | Gene is responsive to IGF-1 [1–3] |
| Cxcl1 | TGAAGCTCCCTTGGTTCAGA<br>AGGTGCCATCAGAGCAGTCT | 3<br>4 | 92 | Gene is responsive to IGF-1 [2, 3] |
| Hspa8 | CATTTGTGTGGTCTCGTCGT<br>CCAACTGCAGGTCCCTTAGA | 1<br>2 | 73 | Gene is responsive to IGF-1 [2, 3] |
| Rgs5 | GCCAGCCAAAATGTGTAAGG<br>TCTGGAGGAGAATTCCCAACT | 1<br>2 | 95 | Gene is responsive to IGF-1 [2, 3] |
| Igfbp3 | CAGGCAGCCTAAGCACCTAC<br>CTTTCCACACTCCCAGCATT | 1<br>2 | 91 | Gene is part of IGF pathway and is responsive to IGF-1 [3] |
| Igf1 | GACCGAGGGGCTTTTACTTC<br>GCAACACTCATCCACAATGC | 3<br>4 | 89 | Gene is part of IGF pathway and is responsive to IGF-1 [3] |
| Serpina7 | AAGTGGCTTCTTGGGCATGT<br>TGATGGTGTGTTCAGCAAGA | 1<br>2 | 92 | Gene is responsive to IGF-1 [3] |
| Vegfa | GCTTGAGTTAAACGAACGTACTTG<br>GAGAGGTCTGGTTCCCGAAA | 4<br>5 | 100 | Gene is responsive to IGF-1 [4, 5] |

**Table 2S.** Wall mounting in the spontaneous alterations test

| Treatment | Sex | 65-week-old | | 98-week-old | |
|---|---|---|---|---|---|
| | | Total tested | Wall mounted | Total tested | Wall mounted |
| Control | M | 25 | 2 | 21 | 0 |
| | F | 24 | 3 | 20 | 1 |
| NVP-ADW742 | M | 25 | 5 | 23 | 1 |
| | F | 25 | 9 | 24 | 5 |
| PPP | M | 22 | 1 | 17 | 0 |
| | F | 25 | 6 | 20 | 2 |

**Table 3S.** Gross necropsy report

| Pathology | Males (group/n) | | | Females (group/n) | | | All (group/n) | | | |
|---|---|---|---|---|---|---|---|---|---|---|
| | Control/25 | NVP-ADW742/25 | PPP/24 | Control/25 | NVP-ADW742/25 | PPP/25 | Control/50 | NVP-ADW742/50 | PPP/50 | Total/149 |
| Seminal vesicle abscess/necrosis | 8 | 8 | 7 | 0 | 0 | 0 | 8 | 8 | 7 | 23 |
| Urinary obstruction | 6 | 1 | 7 | 3 | 0 | 3 | 9 | 1 | 10 | 20 |
| GI bleeding | 0 | 0 | 1 | 1 | 1 | 5 | 1 | 1 | 6 | 8 |
| Heart failure | 3 | 3 | 1 | 5 | 7 | 5 | 8 | 10 | 6 | 24 |
| Liver failure | 7 | 9 | 12 | 4 | 6 | 7 | 11 | 15 | 19 | 45 |
| Kidney failure | 0 | 2 | 1 | 0 | 0 | 0 | 0 | 2 | 1 | 3 |
| GI tumor | 0 | 1 | 0 | 2 | 0 | 1 | 2 | 1 | 1 | 4 |
| Lung tumor | 5 | 7 | 2 | 2 | 1 | 1 | 7 | 8 | 3 | 18 |
| Liver tumor | 8 | 10 | 7 | 2 | 6 | 7 | 10 | 16 | 14 | 40 |
| Seminal vesicle tumor | 1 | 0 | 1 | 0 | 0 | 0 | 1 | 0 | 1 | 2 |
| Pancreas tumor | 4 | 3 | 0 | 2 | 2 | 1 | 6 | 5 | 1 | 12 |
| Adrenal tumor | 0 | 0 | 0 | 2 | 0 | 0 | 2 | 0 | 0 | 2 |
| Kidney tumor | 1 | 0 | 0 | 1 | 1 | 1 | 2 | 1 | 1 | 4 |
| Testis tumor | 0 | 1 | 0 | 0 | 0 | 0 | 0 | 1 | 0 | 1 |
| Anemia | 1 | 0 | 1 | 2 | 1 | 6 | 3 | 1 | 7 | 11 |
| Brain tumor | 0 | 1 | 0 | 4 | 2 | 1 | 4 | 3 | 1 | 8 |
| Ovarian tumor | 0 | 0 | 0 | 1 | 1 | 1 | 1 | 1 | 1 | 3 |
| Vaginal tumor | 0 | 0 | 0 | 0 | 0 | 1 | 0 | 0 | 1 | 1 |
| Spline tumor | 0 | 4 | 1 | 2 | 1 | 3 | 2 | 5 | 4 | 11 |
| Lymphoma | 0 | 0 | 0 | 1 | 1 | 0 | 1 | 1 | 0 | 2 |
| Mouth tumor | 0 | 0 | 0 | 1 | 0 | 0 | 1 | 0 | 0 | 1 |
| Leukemia | 0 | 0 | 0 | 0 | 3 | 4 | 0 | 3 | 4 | 7 |
| Neck tumor | 0 | 0 | 0 | 0 | 0 | 2 | 0 | 0 | 2 | 2 |
| Skin tumor | 0 | 0 | 0 | 1 | 0 | 0 | 1 | 0 | 0 | 1 |
| Tumor unknown | 1 | 0 | 0 | 0 | 1 | 1 | 1 | 1 | 1 | 3 |
| Unknown | 1 | 0 | 0 | 2 | 1 | 0 | 3 | 1 | 0 | 4 |
| Mice with neoplasms | 15 | 18 | 11 | 17 | 18 | 19 | 32 | 36 | 30 | 98 |
| All neoplasms | 21 | 27 | 12 | 23 | 20 | 30 | 44 | 47 | 42 | 133 |

**Table 4S.** Composition of physiological, external, and internal solutions used in hERG channel test

| Reagent | Physiological Solution (mM) | External Solution (mM) | Internal Solution (mM) |
|---|---|---|---|
| NaCl | 140 | 80 | 10 |
| KCl | 4 | 4 | 10 |
| KF | - | - | 110 |
| CaCl2 | 2 | 2 | - |
| MgCl2 | 1 | 1 | - |
| Glucose | 5 | 5 | - |
| NMDG | - | 60 | - |
| HEPES | 10 | 10 | 10 |
| EGTA | - | - | 10 |

**Table 5S.** Sources of liver microsomes for liver microsome stability tests

| Species | Product Information | Vendor |
|---|---|---|
| Human | Cat No. 452117/Lot No. 38298 | Corning-Discovery Labware (Glendale, Arizona) |
| CD-1 Mouse | Cat No. LM-XS-02M/Lot No. TQLK | Rild Research Institute for Liver Disease Co. (Shanghai, China) |

### SUPPLEMENTARY METHODS:

### Identification of NVP-ADW742 drug-likeness properties

#### 1. Water solubility

Aqueous solubility (PBS, pH 7.4) was determined by Eurofins Discovery Services North America, LLC (St. Charles, MO) using a 24-hour shake flask at room temperature following standard protocols [6]. Diethylstilbesterol, disulfiram, metoprolol, phenytoin, rifampicin, and simvastatin were used as reference compounds. A chromatogram of the test compound (200 μM), along with a UV/VIS spectrum with labeled absorbance maxima, was generated. Aqueous solubility (μM) was determined by comparing the peak area of the principal peak in a calibration standard (200 μM) containing organic solvent (methanol/water, 60/40, v/v) with the peak area of the corresponding peak in a buffer sample.

#### 2. Igf1R dissociation constant (Kd)

Igf1R Kd was measured using IGF1R Human RTK Kinase KINOMEscan KdELECT Binding LeadHunter Assay (Cat. # 87-0007-1198, Eurofins Discovery Services North America, LLC, St Charles, MO). The assay is based on a competitive binding assay that quantitatively measures a compound's ability to compete with an immobilized, active-site-directed ligand [7]. The assay is performed by combining three components: DNA-tagged kinase; immobilized ligand; and a test compound. The ability of the test compound to compete with the immobilized ligand is measured via quantitative PCR of the DNA tag. Kd was determined using an 11-point 3-fold compound dilution series with three 0.9% DMSO control points.

### 3. Igf1R half-maximal inhibitory concentration (IC50) in human cell-line

IC50 was determined using Cell-Based Antagonist Functional LeadHunter Assays for human Igf1R (Cat. # 86-0006P-2755AN, Eurofins Discovery Services North America, LLC, St Charles, MO). This assay HEK293 utilizes PathHunter Receptor Tyrosine Kinase (RTK) functional cell lines stably transfected to co-express a ProLink™ (PK) tagged RTK (Igf1R in our case) and an enzyme acceptor (EA) tagged SH2 domain [8]. Activation of the RTK-PK induces receptor dimerization, leading to SH2-EA recruitment, and forcing complementation of the two β-galactosidase enzyme fragments (EA and PK). The resulting functional enzyme hydrolyzes the substrate to generate a chemiluminescent signal.

### 4. Insulin receptor (IR) IC50

IC50 was determined using IR Human Kinase Enzymatic Radiometric LeadHunter Assay (Cat. #174990, Eurofins Panlabs Discovery Services Taiwan Ltd), which utilizes human recombinant IR. NVP-ADW742 was preincubated with 0.067 µg/ml IR for 15 minutes at 37°C in modified HEPES buffer pH 7.4. The reaction was initiated by the addition of 0.2 mg/ml poly(Glu:Tyr), 10 µM ATP containing 0.25 µCi [γ32P]ATP for 30 minutes, and terminated by further addition of 3% H3PO4. Poly(Glu:Tyr) is a synthetic substrate for measuring the tyrosine kinase activity, where active kinase transfers phosphate to the substrate. Radioactivity of [32P]Poly(Glu:Tyr) was counted to identify IR activity with and without NVP-ADW742. IC50 values were determined by a non-linear, least squares regression analysis using MathIQTM (ID Business Solutions Ltd., UK).

### 5. HepG2 cell viability

The goal of the HepG2 cell viability assay is to compare a compound's effect on both glucose and galactose-grown cells to identify mitochondrial toxicants. If a compound is toxic in galactose-grown cells, but not in glucose, then it suggests that the compound impairs mitochondrial function. However, if a compound demonstrates toxicity in both glucose and galactose-grown cells, then it suggests that while there may be mitochondrial dysfunction occurring, it is most likely secondary to other effects because it is impairing glycolysis as well. HepG2 viability assay was done by WuXi AppTec Ltd (Kowloon, Hong Kong) following standard protocols. HepG2 cells were seeded in 30 µl Glucose medium or Galactose (90% DMEM, 10% FBS, 5 mM HEPES(1M), 2mM L-Glutamine (200 mM), 1x Penicillin-Streptomycin (100x)) and incubated overnight at 37 ℃ with 5% CO2. Cells were then treated with 30 µl of NVP-ADW742 for 24 hours (Final DMSO 0.5%--2.8%), and 20 µl of CTG (Promega, Cat# G7573) was added to each well. Following 10 min incubation, ATP concentration was determined by reading luminescence in a Perkin Elmer Envision plate reader.

### 6. hERG channel test on SyncroPatch

A hERG assay measures if a compound inhibits the hERG potassium channel, which can cause dangerous cardiac arrhythmias (cardiotoxicity). The hERG channel assay was performed by WuXi AppTec Ltd (Kowloon, Hong Kong) using an automated patch-clamp system, SyncroPatch, to measure the actual potassium current through the channel in cultured cells. CHO cells stably expressing hERG potassium channels from Sophion Biosciences were used for this test. The cells were cultured in CHO hERG Culture Medium (Ham's F12 – 500 ml, FBS – 50 ml, G418/Geneticin – 1 ml, and Hygromycin B – 1 ml) in a humidified and air-controlled (5% CO2) incubator at 37 °C. 75% confluent cells were harvested two days after plating using TrypLE and resuspended in the physiological solution at room temperature. For the electrophysiological recordings, the following solutions were used (Table 4S). NVP-ADW742 was dissolved in 100% DMSO to obtain a stock solution for different test concentrations. Then the stock solution was diluted into the external solution to achieve final concentrations for testing. Final DMSO concentration in the extracellular solution was not more than 0.30% for the test compounds. Voltage command protocol is shown in Fig. 3S: from the holding potential of -80 mV, the voltage was first stepped to -50 mV for 80 ms for leak subtraction and then stepped to +20 mV for 4800 ms to open hERG channels. After that, the voltage was stepped back down to -50 mV for 5000 ms, causing a "rebound" or tail current, which was measured and collected for data analysis. Finally, the voltage was stepped back to the holding potential (-80 mV, 1000 ms). This voltage command protocol was repeated every 20000 msec. This command protocol was performed continuously during the test (vehicle control and test compound). Five concentrations (0.30 μM, 1.00 μM, 3.00 μM, 10.00 μM and 30.00 μM) were tested for each compound. Minimum 2 replicates per concentration were obtained.

Data analysis was carried out using DataControl, Excel 2013 (Microsoft), and GraphPad Prism 5.0. Within each well recording, the percent of control values was calculated for each test compound concentration, current response based on peak current in the presence of reference control (current response/ peak current) ×100%. The Dose-Response curves were fit to the standard Hill equation as shown below:

Ipost cpd/Ipre cpd=Bottom+(Top-Bottom)/(1+10^((LogIC50-X)*HillSlope))

Where X is the logarithm of concentration, Ipost cpd/Ipre cpd is the normalized peak current amplitude, Top is 1, and Bottom is equal to 0.

### 7. Liver microsome stability

A microsomal stability test was performed by WuXi AppTec Ltd (Kowloon, Hong Kong) using standard protocols to measure NVP-ADW742 metabolic degradation rate by liver microsomes, which is predictive of in vivo metabolic stability and hepatic clearance [9]. In short, compound working solutions (CWS) of NVP-ADW742 and control compounds (testosterone, diclofenac, and propafenone) were prepared by diluting 10 mM stock solutions in dimethyl sulfoxide (DMSO) to 100 μM with acetonitrile (ACN). 10 mM working solution of β-Nicotinamide adenine dinucleotide phosphate reduced form, tetrasodium salt (NADPH·4Na, Cat. # BT04, Bontac Bio-Engineering LTD, Shenzen, China) was prepared in 10 mM MgCl2 solution. To prepare microsome working solutions (MWS), microsomes (Table 5S) were diluted to a concentration of 0.56 mg/mL using a 100 mM potassium phosphate buffer (PB). ACN containing 250 nM tolbutamide and 250 nM labetalol as internal standards (IS) was used as a stop solution.

MWS (445 μL) was transferred into pre-warmed 'Incubation' plates T60 and NCF60, followed by a 10 min incubation at 37°C. Next, 54 μL of MWS was transferred to a Blank60 plate, followed by the addition of 6 μL NADPH cofactor and 180 μL of stop solution into each well. CWS (5 μL) was added to the 'incubation' plates (T60 and NCF60) containing microsomes. 50 μL of PB buffer was added to NCF60 plate and the plate was incubated at 37°C for 60 min. Stop solution (180 μL) and NADPH working solution (6 μL) were added to the T0 plate. Then, a mixture (54 μL) was removed from the 'Incubation' plate T60 and transferred to the T0 plate. NADPH working solution (44 μL) was added to the T60 plate, followed by a 60 min incubation at 37°C. At 5, 15, 30, 45, and 60 min, 60 μL of each sample at each time point was transferred to a well containing 180 μL of stop solution, followed by mixing. All sampling plates were shaken for 10 min, then centrifuged at 3220 ×g for 20 min at 4°C. The supernatant (80 μL) was transferred into 240 μL of pure water and mixed using a plate shaker for 10 min. Each bioanalysis plate was sealed and shaken for 10 min before LC-MS/MS analysis (see Supplemental method 11). Half-life was calculated using first-order kinetics equations.

### 8. Bi-directional permeability across the Caco-2 cell monolayer

Bi-directional Caco-2 cell permeability assays were performed by WuXi AppTec Ltd (Kowloon, Hong Kong) to predict oral absorption and identify active efflux mechanisms, by measuring the rate of compound transport across a confluent monolayer in both apical-to-basolateral (A→B, absorption) and basolateral-to-apical (B→A, secretion) directions. Human Caco-2 cells purchased from ATCC were seeded onto 0.4 μm pore polycarbonate membranes (PC) in 96-well Corning Insert plates at 3.5 x 104 cells/cm2, and medium was refreshed every 4~5 days until the 21st to 28th day for confluent cell monolayer formation.

NVP-ADW742 was tested at 2.00 µM bi-directionally in duplicate. Nadolol was used as a low permeability marker; metoprolol as a high permeability marker; and digoxin as a positive control for P-glycoprotein (P-gp) substrate. Digoxin was tested at 10.0 µM bi-directionally in duplicate, while nadolol and metoprolol were tested at 2.00 µM in A to B direction in duplicate. Final DMSO concentration was adjusted to less than 1.0%. The plate was incubated for 2 hours in CO2 incubator at 37.0°C, with 5% CO2 at saturated humidity without shaking. After being mixed with the stop solution, all samples were centrifuged at 3220xg for 10 minutes. Concentrations of test and control compounds in all samples were semiquantitatively determined by LC-MS/MS methodologies, using the area ratio of analyte/internal standard. After the transport assay, the lucifer yellow rejection assay was applied to determine the Caco-2 cell monolayer integrity. The transport buffer in the study was HBSS (Hanks Balanced Salt Solution) containing 10.0 mM HEPES (2-[4-(2-Hydroxyethyl)-1-piperazinyl] ethanesulfonic acid) at pH 7.40 ± 0.05.

The apparent permeability coefficient $P_{app}$ (cm/s) was calculated using the equation:

$$P_{app} = (dC_r/dt) \times V_r / (A \times C_0)$$

Where $dC_r/dt$ is the cumulative concentration of compound in the receiver chamber as a function of time; $V_r$ is the solution volume in the receiver chamber (0.0750 mL on the apical side, 0.250 mL on the basolateral side); A is the surface area for the transport, i.e. 0.143 cm2 for the area of the monolayer; $C_0$ is the initial concentration in the donor chamber.

The efflux ratio was calculated using the equation:

$$\text{Efflux Ratio} = P_{app}\ (B\text{-}A) / P_{app}\ (A\text{-}B)$$

Percent recovery was calculated using the equation:

$$\%\text{Solution Recovery} = 100 \times [(V_r \times C_r) + (V_d \times C_d)] / (V_d \times C_0)$$

Where $V_d$ is the volume in the donor chambers (0.0750 mL on the apical side, 0.250 mL on the basolateral side); $C_d$ and $C_r$ are the final concentrations of transport compound in donor and receiver chambers, respectively.

#### 9. In vivo pharmacokinetics (PK)

Male 6-9-week-old C57BL/6 mice were purchased from Shanghai SIPPR-Bk Lab Animal Co., Ltd and used for PK study conducted by WuXi AppTec Ltd (Kowloon, Hong Kong). Three animals were used in the intravenous (IV) protocol, and three animals were used in the per-oral (PO) protocol. Before being placed on study, animals were acclimated to the test facility for at least 3 days and checked for their general health by veterinary staff or other authorized personnel at the end of the acclimation period. Animals were group-housed up to four animals/cage in polysulfone cages with certified corncob bedding during acclimation and study period. Environment controls were set to maintain a temperature range of 20-26℃, a relative humidity range of 40 to 70%, and a 12-hour light/12-hour dark cycle. Certified rodent diet and water were provided to all animals ad libitum.

1 mg/mL NVP-ADW742 sterile solution in NMP/PEG300 (10:90) was prepared and filtered through a 0.22 µm filter before dosing. Animals were dosed within four hours after the formulation was prepared._The formulation sample was then removed from the formulation solutions, transferred into 1.5 mL polypropylene microcentrifuge tubes, and used for dose validation by LC-MS/MS. An LC-MS/MS method has been developed with a calibration curve consisting of 6 calibration standards (see Supplemental method 11). Acceptance criteria for an analytical run were that at least 5 of 6 calibration standards should be within ±20% of nominal values. The nominal IV bolus dose was 3 mg/kg body weight, and the administered dose was 2.82 mg/kg body weight. The nominal PO dose was 10 mg/kg body weight, and the administered dose was 9.41 mg/kg body weight.

For PO dosing, the dose formulation was administered via oral gavage following the facility's SOPs. For IV dosing, the dose formulation was administered via the tail vein. The dose volume was determined by the animals' body weight collected on the morning of dosing day. For IV administration, animals were fasted overnight before the treatment.

Blood collection (about 0.025 mL per time point) was performed via saphenous vein into commercial microcentrifuge tubes containing K2-EDTA as an anti-coagulant and placed on wet ice until centrifugation. For the PO dosing experiment, blood was collected following 0.25, 0.5, 1, 2, 4, 8, and 24 hours post-dosing. For the IV dosing experiment, blood was collected following 0.083, 0.25, 0.5, 1, 2, 4, 8, and 24 hours post-dosing. Samples were centrifuged (3200×g for 10 minutes at 2 to 8°C) within one hour of collection. The plasma samples were transferred into labeled polypropylene micro-centrifuge tubes and stored frozen in a freezer at -60°C or lower until bio-analysis. All animals were observed at dosing and each scheduled collection, and all abnormalities were recorded. A bioanalytical method has been developed for LC-MS/MS quantitative determination of NVP-ADW742 in biological matrix (see Supplemental method 11). Plasma concentration versus time data were plotted and analyzed by non-compartmental approaches using the Phoenix WinNonlin 8.3.5 software program.

#### 10. Mouse Brain and Plasma Protein Binding

Equilibrium dialysis method was used by WuXi AppTec Ltd (Kowloon, Hong Kong) to measure drug-protein binding. This method uses a semi-permeable membrane to separate protein-bound drug from the free (unbound) fraction. The unbound drug passes through the membrane until equilibrium is reached, allowing for accurate calculation of binding percentages in plasma or tissue. All analyses were done in triplicate.

HT-Dialysis plate (Model HTD 96 b, Cat # 1006) and the dialysis membrane (molecular weight cut off 12 ~ 14 kDa, Cat # 1101) were purchased from HT Dialysis LLC (Gales Ferry, CT). BupHTM Phosphate Buffered Saline Pack, supplied by Thermo Fisher Scientific under Product # 28372, contained 0.1 M sodium phosphate and 0.15 M sodium chloride with a pH of 7.2, when the pouch contents were dissolved in a final volume of 500 mL deionized water. Before use, the pH value was adjusted to pH 7.4 ± 0.1 using 1% phosphoric acid or 1 N sodium hydroxide.

On the day of the experiment, mouse plasma was thawed by running under cold tap water and centrifuged at 3220 ×g for 5 minutes to remove any clots. The pH value of the resulting plasma was measured and adjusted to pH 7.4 ± 0.1 using 1% phosphoric acid or 1 N sodium hydroxide, if required. Mouse brain homogenate samples were thawed in a water bath at room temperature and then incubated at 37°C for 10 minutes before use.

Dialysis membrane strips (containing 2 membranes) were soaked in ultra-pure water at room temperature for approximately 1 hour. Then the two membranes were separated and soaked in ethanol: water (20: 80, v: v) for approximately 20 minutes. Prior to use, the membranes were rinsed and soaked for 20 minutes in ultra-pure water. Working solutions (400 μM) of the test compound (NVP-ADW742) and control compound (warfarin for plasma protein binding or propranolol for brain homogenate protein binding) were prepared by diluting 4 μL of the stock solution with 96 μL DMSO. Loading matrix solutions (2 μM) of both compounds were prepared by diluting 3 μL of the working solutions with 597 μL of blank matrix and were mixed thoroughly. The stop solution was 250 nM tolbutamide and 250 nM labetalol in acetonitrile.

Aliquots of 50 μL loading matrix containing test or control compound were transferred in triplicate to the Sample Collection Plate. The samples were matched with opposite blank PBS to obtain a final volume of 100 μL with a volume ratio of matrix: PBS at 1: 1 (v: v) in each well immediately. The stop solution (500 μL) was added to these T0 samples of test compound and control compound. The plate was sealed and shaken at 800 rpm for 10 minutes. Then these T0 samples were stored at 2-8°C pending further process, along with other post-dialysis samples.

An aliquot of 100 µL of the loading matrix containing test or control compounds was transferred to the donor side of each dialysis well in triplicate, and 100 µL of PBS was loaded to the receiver side of the well. Then the plate was rotated at 100 rpm in a humidified incubator with 5% carbon dioxide at 37°C for 4 hours. At the end of the dialysis, aliquots of 50 µL samples from the PBS (receiver) side and the matrix (donor) side of the dialysis device were taken into new 96-well plates (Sample Collection Plates). An equal volume of opposite blank PBS or matrix in each sample was added to reach a final volume of 100 µL with a volume ratio of matrix: PBS at 1: 1 (v: v) in each well. Stop solution (500 µL) was added to these samples. The mixture was vortexed and centrifuged at 4000 rpm for about 20 minutes. An aliquot of 100 µL of supernatant of all samples was then removed for LC-MS/MS analysis. The single blank sample was prepared by transferring 50 µL of blank matrix to a 96-well plate and adding 50 µL of blank PBS to each well. The matrix-matched samples were processed by adding 500 µL of stop solution containing internal standards, following the same sample processing method as the other samples. Protein binding was calculated using the following equations:

% Unbound = 100 × F / T,

% Bound = 100 - % Unbound,

% Recovery = 100 × (F + T) / T0,

where

F = the peak area ratio of compound and internal standard on the receiver side of the membrane after 4 hours of incubation,

T = the peak area ratio of compound and internal standard on the donor side of the membrane after 4 hours of incubation,

T0 = the peak area ratio of compound and internal standard at time zero.

#### 11. PK in plasma and brain following a single IV injection

Male 6-9-week-old C57BL/6 mice were purchased from Shanghai SIPPR-Bk Lab Animal Co., Ltd and used for a PK study conducted by WuXi AppTec Ltd (Kowloon, Hong Kong). Nine animals were used in this study, three per time point (see below). Before being placed on study, animals were acclimated to the test facility for at least 3 days and checked for their general health by veterinary staff or other authorized personnel at the end of the acclimation period. Animals were group-housed up to four animals/cage in polysulfone cages with certified corncob bedding during acclimation and study period. Environment controls were set to maintain a temperature range of 20-26℃, a relative humidity range of 40 to 70%, and a 12-hour light/12-hour dark cycle. Certified rodent diet and water were provided to all animals ad libitum.

1 mg/mL NVP-ADW742 sterile solution in NMP/PEG300 (10:90) was prepared and filtered through a 0.22 µm filter before dosing. Animals were dosed within four hours after the formulation was prepared._The formulation sample was then removed from the formulation solutions, transferred into 1.5 mL polypropylene microcentrifuge tubes, and used for dose validation by LC-MS/MS. The nominal IV bolus dose was 3 mg/kg body weight, and the administered dose was 2.94 mg/kg body weight.

For IV dosing, the dose formulation was administered via the tail vein. The dose volume was determined by the animals' body weight collected on the morning of dosing day. For IV administration, animals were fasted overnight before the treatment.

Blood collection (about 0.025 mL per time point) was performed via saphenous vein into commercial microcentrifuge tubes containing K2-EDTA as an anti-coagulant and placed on wet ice until centrifugation. After blood collection, animals were euthanized by $CO_2$ inhalation at 0.083, 0.25, and 1 hours post-dose. After euthanasia, each animal underwent whole-body perfusion with saline through the heart, and then the brain tissue was collected from each animal. After collection, the brain tissue was washed with cold saline, wiped dry, and weighed.

Blood samples were centrifuged (3200×g for 10 minutes at 2 to 8°C) within one hour of collection. The plasma samples were transferred into labeled polypropylene micro-centrifuge tubes and stored frozen in a freezer set to maintain -60°C or lower until bio-analysis. Brain tissue was homogenized using homogenizing buffer (15 mM PBS (pH7.4):MeOH=2:1) at a ratio of 1:10 (1 g tissue with 10 mL buffer, the dilution ratio is 11). The tissue homogenate was kept at -60℃ or lower until LC-MS/MS analysis.

LC-MS/MS quantitative determination of NVP-ADW742 in biological matrix was done as described below (see Supplemental method 11). Plasma and brain concentration versus time data were plotted and analyzed by non-compartmental approaches using the Phoenix WinNonlin 8.3.5 software program.

#### 12. Analytical Method for NVP-ADW742 quantification in mouse plasma

Instrument: LC-MS/MS-CM_API5000

Matrix: Male C57BL/6J Mouse Plasma EDTA-2K

Analyte: NVP-ADW742

Internal standard(s): IS1:6 in 1 internal standard in ACN (Labetalol & tolbutamide & Verapamil & dexamethasone & glyburide & Celecoxib 100 ng/mL for each)

MS conditions: ESI:Positive

MRM detection

NVP-ADW742:[M+H]+m/z: 454.40/383.20 Da

Verapamil:[M+H]+m/z: 455.20/164.90 Da

UPLC conditions Mobile Phase:

Mobile Phase A:0.1% FA & 2mM NH4OAc in water/ACN (v:v, 95:5)

Mobile Phase B:0.1% FA & 2mM NH4OAc in water/ACN (v:v, 5:95)

| Time (min) | Mobile Phase B (%) |
|---|---|
| Initial | 5 |
| 0.2 | 5 |
| 1.1 | 100 |
| 1.4 | 100 |
| 1.41 | 5 |
| 1.6 | 5 |

Column: ACQUITY UPLC BEH C18 1.7 μm 2.1 × 50 mm Column

Column temperature: 45.0 C

Flow rate: 0.6 mL/min

Retention time:

NVP-ADW742: 0.965 min

Verapamil: 1.07 min

Sample preparation:

1) Aliquots of 3 μL of the unknown sample, calibration standard, quality control, dilution quality control, single blank, and double blank samples were added to the 96-well plate.

2) Each sample (except the double blank) was quenched with 60 µL of IS1 (the double blank sample was quenched with 60 µL of ACN), and then the mixture was vortex-mixed for 10 min at 800 rpm and centrifuged for 15 min at 3220 × g, 4°C.

3) An aliquot of 55 µL supernatant was transferred to another clean 96-well plate and centrifuged for 5 min at 3220 × g, 4°C, then the supernatant was directly injected for LC-MS/MS analysis.

Dilution procedure description:

An aliquot of 2 µL of the unknown sample was added to 18 µL blank matrix.

Calibration curve:

1.00-3000 ng/mL for NVP-ADW742 in Male C57BL/6J Mouse Plasma EDTA-2K

Other bioanalytical guidance:

Follow DMPK-Lab-1201-005 Standard Operating Procedure for LC-MS Based Quantitative Analysis
